# Does colour impact attention towards 2D images in geckos?

**DOI:** 10.1101/2021.02.03.429594

**Authors:** Nathan Katlein, Miranda Ray, Anna Wilkinson, Julien Claude, Maria Kiskowski, Bin Wang, Scott Glaberman, Ylenia Chiari

**Affiliations:** University of South Alabama, Department of Biology, Mobile, AL 36688, USA; University of Lincoln, School of Life Sciences, Lincoln, LN6 7DL, UK; Institut des Sciences de l’Evolution de Montpellier, UMR UM/CNRS/IRD/EPHE, 2, Pl. E. Bataillon, 34095 Montpellier, France; University of South Alabama, Department of Mathematics and Statistics, Mobile, AL 36688, USA; George Mason University, Department of Environmental Science and Policy, Fairfax, VA 22030, USA; George Mason University, Department of Biology, Fairfax, VA 22030, USA

**Keywords:** 2D images, Behaviour, Familiar object, Habituation, Image perception, Novel object, Reptile, Vision

## Abstract

Animals are exposed to different visual stimuli that influence how they perceive and interact with their environment. Visual information such as shape and colour can help the animal detect, discriminate and make appropriate behavioural decisions for mate selection, communication, camouflage, and foraging. In all major vertebrate groups, it has been shown that certain species can discriminate and prefer certain colours and that colours may increase the response to a stimulus. However, because colour is often studied together with other potentially confounding factors, it is still unclear to what extent colour discrimination plays a crucial role in the perception of and attention towards biologically relevant and irrelevant stimuli. To address these questions in reptiles, we assessed the response of three gecko species *Correlophus ciliatus, Eublepharis macularius*, and *Phelsuma laticauda* to familiar and novel 2D images in colour or grayscale. We found that while all species responded more often to the novel than to the familiar images, colour information did not influence object discrimination. We also found that the duration of interaction with images was significantly longer for the diurnal species, *P. laticauda*, than for the two nocturnal species, but this was independent from colouration. Finally, no differences among sexes were observed within or across species. Our results indicate that geckos discriminate between 2D images of different content independent of colouration, suggesting that colouration does not increase detectability or intensity of the response. These results are essential for uncovering which visual stimuli produce a response in animals and furthering our understanding of how animals use colouration and colour vision.

## Introduction

Animals are confronted with a multitude of visual stimuli that affect their interactions with the environment (Dall *et al*., 2005). Although a stimulus with a single component (e.g., animal shape) may be sufficient to elicit a response, many species and some specific functions may integrate two or more components (e.g., shape and colour) to increase the detectability of the stimulus by the receiver (Grether, Kolluru & Nersissian, 2004) and the magnitude of the response (Stevens, 2013). Colour variation and colour vision are among the most studied visual stimuli for their role in adaptation, as they are used for communication, including mate choice, sexual selection, and intrasexual competition, foraging, camouflage and background matching (e.g., Hubbard *et al*., 2010; Olsson, Stuart-Fox & Ballena, 2013; Cuthill *et al*., 2017; Zambre & Thaker, 2017; Caro & Mallarino, 2020). Representatives of all major vertebrate groups have been shown to be able to detect colour and colour contrast and even show preference for specific colours (e.g., Sherwin & Glen, 2003; Roth & Kelber, 2004; Hansen, Beer & Müller, 2006; Luchiari & Pironhen, 2008; Svádová *et al*., 2009; Passos, Mello & Young, 2014; Maia *et al*., 2017; Yovanovich *et al*., 2017; Smithers *et al*., 2018), indicating that colouration alone can elicit a response in animals. Furthermore, colouration in combination with scent, sound or motion may modulate the detection of the stimulus (for review see Hubbard *et al*., 2010; Olsson *et al*., 2013; Cuthill *et al*., 2017; Caro & Mallarino 2020). Yet, the impact of colouration alone on perception of biologically relevant vs irrelevant (unfamiliar) stimuli, and the magnitude of that impact is still relatively unclear, especially for non-human vertebrates (e.g., Delorme, Richard & Fabre-Thorpe, 2000; Tanaka, Weiskopf & Williams, 2001; Delorme, Richard & Fabre-Thorpe, 2010; Klomp *et al*., 2017; Schwedhelm, Baldauf & Treue, 2020). Such research will further our knowledge on how animals use colouration and colour vision, especially depending if they are familiar or not with the observed stimulus.

In reptiles, research on visual stimuli has generally focused on the role of colouration in an ecological and evolutionary context (but see Wilkinson, Mueller-Paul & Huber, 2013; Frohnwieser *et al*., 2017), especially in combination with other stimuli such as movement, scent, or shape information (e.g., LeBas & Marshall, 2000; Klomp *et al*., 2017; Ossip-Drahos *et al*., 2018; Kabir, Radhika & Thaker, 2019; Dollion *et al*., 2020). It is therefore difficult to tease apart the role that colour alone plays in the process of object discrimination, especially in the context of biologically relevant vs irrelevant images. For example, Frohnwieser *et al*. (2017) found that colour information was not necessary for lizards (*Pogona vitticeps*) to perceive an object shaped like a lizard as a lizard. To our knowledge this is the only work that has investigated this issue and did so using only one tested species. Due to the lack of similar studies also on other lizard species, it remains unclear is the influence that colouration has on 2D object perception when other confounding factors are removed (see Kleiber *et al*., 2021 for a similar approach in fish). Here, study how three species of geckos (order Squamata) –*Phelsuma laticauda, Correlophus ciliatus* and *Eublepharis macularius* – interact with images with differing colour content. Specifically, we use an object familiar to the geckos (an image of a conspecific gecko) and a randomly chosen unfamiliar object (an image of a car), both in colour and grayscale. We then record whether or not the geckos interact with the coloured and grayscale images and measure the duration of each interaction when occurring. The three test species differed in their habitat use (terrestrial vs. arboreal), main time of activity (nocturnal vs. diurnal), origin (captive bred vs. wild caught), body colour, and colour pattern, which may influence how they respond to colour information.

Geckos have largely been overlooked among reptiles in studies focusing on responses to visual stimuli (but see Kabir *et al*., 2019), possibly because most, but not all (Gamble *et al*., 2015), genera in this group are nocturnal. All geckos evolved from diurnal lizards and have retained eyes comprised of cone-derived photoreceptors that are used for colour vision even in low light conditions (Röll, 2000; Roth & Kelber, 2004; Pinto, Nielsen & Gamble, 2019). Geckos also possess tetrachromatic colour vision with cones sensitive to ultraviolet (UV), blue, and green light (Röll, 2000; Roth & Kelber, 2004). Colour and colour pattern variation is extremely prominent both within and between species (Allen *et al*., 2020), including sexual dimorphism and sexual dichromatism (Lukas & Frynta, 2002; Marcellini, 1977), suggesting that these characters could be used in communication among individuals and species, and for sexual selection. However, despite the large variation in colour and colour pattern observed in geckos and the diverse biology of these animals, little is known about how colour information is perceived and, to our knowledge, on how colour is used to improve perception.

Although all species used in this work can see colour, we expect the brightly coloured diurnal species (*P. laticauda*) to rely more on colour information – potentially used in communication among conspecifics – and therefore show greater responses to coloured images than nocturnal species. Furthermore, we also anticipate a dissimilar response between the gecko images shown in their natural vs. grayscale colour, as the gecko is both familiar and biologically relevant. The use of the unfamiliar image (car) allows us to examine impact of colour on an object that was not biologically relevant. As such, we do not expect to find any difference between the coloured vs. grayscale images of the unfamiliar object. The data collected in this work will improve our understanding of how colour is used as a visual stimulus in geckos, with implications for their foraging, mating, and communication.

## Materials and Methods

### Ethical note

All capture, handling and experimental protocols were approved by the University of South Alabama IACUC committee (protocol #993866). Experiments were carried out to minimize stress and disturbance to animals and in accordance with relevant guidelines and regulations.

### Experimental Set-Up and Testing Procedure

Seven to eight geckos per species were used in experiments (Table 1). Tests were run in an arena (30.5 cm x 61 cm x 20.3 cm) constructed of clear plastic and silicone. In order to test how geckos interact with coloured visual stimuli, we used printed images that were presented only on one side of the testing arena. All experiments were recorded using two cameras (Supplementary Material Fig. S1). To test if geckos interact differently with a coloured vs. non-coloured object, we presented pictures of each object in their natural colour or in grayscale. Grayscale was used as an alternative to natural colour to eliminate the hue and saturation and leave just the brightness and intensity of each pixel in a given image (see Supplementary Material for details). We therefore refer to colour in this work as to the chromatic components of colouration, since brightness was not modified. To understand if colour impacts object perception, especially if biologically relevant vs irrelevant, we selected two distinct objects: (1) an image of each gecko species and (2) an image of a car. A car was chosen as a random unfamiliar irrelevant object as most likely geckos have never seen it before (for an example of the use of randomly selected objects never seen by the tested species see also Kleiber *et al*., 2021). For each object type (here called image type), two different images of the same type (e.g., two individual geckos) were used to ensure that the responses obtained were not due to the specific image shown but rather the type of image itself (Supplementary Material Fig. S2). Within a single week, geckos were shown only one image type in both colour and grayscale and each trial lasted for 10 minutes for each image (Supplementary Material Fig. S3). In our experimental set up, we tested for differences in responses based on species, sex, image colour, image type, different images of the same type, image order based on colour, as well as habituation or memory to the experimental setup (Supplementary Material Table S1). Complete details on the captivity conditions, facilities, and experimental set up can be found in Supplementary Material.

**Table 1:**
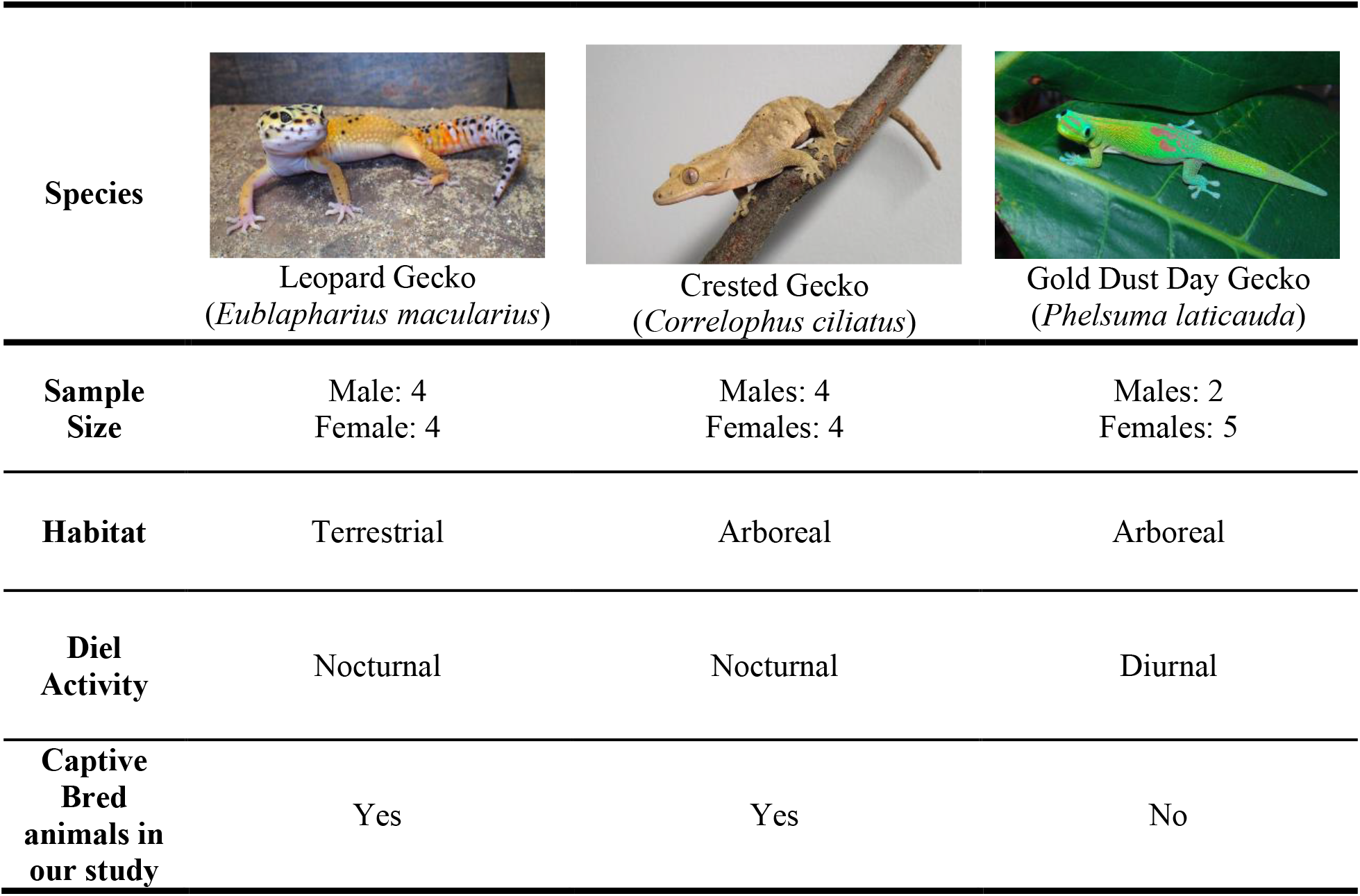
Characteristics of species used in the experiment (IUCN, 2018). Image credit: Tony Gamble.

### Data Collection and Statistical Analyses

We considered geckos to be interacting with an image (called “response” here) if they looked at, touched, or licked the images during each experiment using the following definitions: *touching* – when a gecko’s tongue (licking) or snout was touching the image; *looking* – when a gecko’s head and eyes were directed directly towards the image. Both behaviours, *touching* and *looking*, were considered as “response”. Gecko were considered *not looking* (defined as absence of a response) when a gecko had faced the image and was aware of it but after that its head and part of its body are not directed towards the image so that it is not looking at it (Supplementary Material Fig. S4, see also Supplementary Material for additional details on these categories and using a more conservative approach on what we considered a “response”).

We analysed the influence of each variable and their interactions on the presence/absence of a response to the image and on the proportion of time of response (defined here as *PLT*) only for geckos that saw the images. Presence or absence of response (*PLT*>0 vs. *PLT*=0, respectively) were coded as geckos that saw the image, moved towards it, touched it, or licked it (*PLT*>0; *touching* and *looking*), versus geckos that saw the image but turned their head away and ignored it (*PLT*=0; *not looking*, see detailed definition above and in Supplementary Material). Analyses were run in R (version 3.5.3, R Core Team, 2017) using the “*glmer*” function from the package “*lme4*” (Bates, Maechler & Bolker, 2012). To study the influence of each explanatory variable on *PLT*, we used a generalized linear mixed effects model, with the variables of interest as fixed effects and individual ID as a random variable for random effects. Individuals were considered as a random factor because the same individuals were used throughout multiple experiments (i.e., different image type, two images of the same type, and colour versus non-colour images) and because we were interested in inter-individual variation in response. We tested the effects of each explanatory variable by performing model comparison using chi-square tests (Supplementary Material Table S1 for description of each variable). Wald tests were then used to test whether the coefficients were significantly different from zero in the coefficient table of the fitted mixed effects models. Tuckey’s method was used to adjust the p-values for multiple comparisons.

We first tested for variation between response (PLT>0) versus non-response (PLT=0) by fitting a mixed effects logistic regression model with binomial distribution. Successively, we use a generalized linear mixed effects model to compare the mean PLTs with data only for PLT>0 to see which factors affected the response of the geckos. Different distributions were checked for PLT>0 data. The Gamma distribution was used in the mixed effects models as it fits the PLT>0 data better than the normal, log-normal and Poisson distributions. For each dataset, we first fitted a full model with all first-order terms of all covariates, and then selected the best model using backward methods. Analyses were re-run after removing non-significant effects in order to make sure that results were the same irrespective of the order of effect removal (results are not reported here because analyses were always consistent). Interaction effects were evaluated among covariates of interest – i.e., colour and species, species and image type, colour and image type, image type and image order, day and image type, sex and colour, sex and image type. Interaction effects were tested one at the time. The R package “*effects*” (Fox, 2003; Fox & Weisberg, 2018) was used to gain insights into the differences among different levels of the factors of interests. A full description of the experimental procedure and variables tested are described in Supplementary Material. Analyses were also repeated using “*touching*” data only (touching vs. all the rest of the responses and then only for *touching*>0, considering for how long the animal touched the image).

## Results

### 3.1 Within species variation in PLT

For *E. macularius*, 128 videos were collected; in 7.8% of these videos, the geckos never saw the image shown; of the remaining videos, in 11.9% the geckos saw the images and did not respond (*PLT*=0) and in 88.1% they responded (*PLT*>0). For *C. ciliatus*, 124 videos were collected; in 28.2% of these videos, the geckos never saw the image shown; of the remaining videos, in 24.7% the geckos saw the image and did not respond (*PLT*=0) and in 75.3% the geckos responded (*PLT*>0). Lastly, for *P. laticauda*, 97 videos were collected; in 10.3% of these videos the geckos never saw the image shown; of the remaining videos, in 24.1% the geckos saw the images and did not respond (*PLT*=0) and in 75.9% they responded (*PLT*>0). Overall, across all the species, we obtained more responses (*PLT*>0) than non-responses (*PLT*=0) to the images. Detailed summary statistics for *PLT* measures can be found in Supplementary Materials Table S2. *Touching*>0 corresponds to 106 entries in total, representing 44.73% of the data. Frequency and distribution of PLT on log transformed and raw data for each species can be found in Supplementary Material Fig. S5.

### 3.2 Non-response (PLT=0) vs. response (PLT>0) analysis

Overall, geckos had a *PLT*=0 (non-response) in 57 out of 294 (19.4%) experiments in which the geckos saw the images. Although *E. macularius* interacted more with images than the other two species, differences were not significant among species (*p*>0.05). Table 2 shows the influence of each variable from a fitted generalized linear effects model. Geckos reacted differently most prominently based on image type (p=0.003), but also image order (p=0.010), test room (p=0.037) and experimental day (p=0.041) (Table 2a). Gecko did not respond differently to coloured versus grayscale images (Table 2).

**Table 2:**
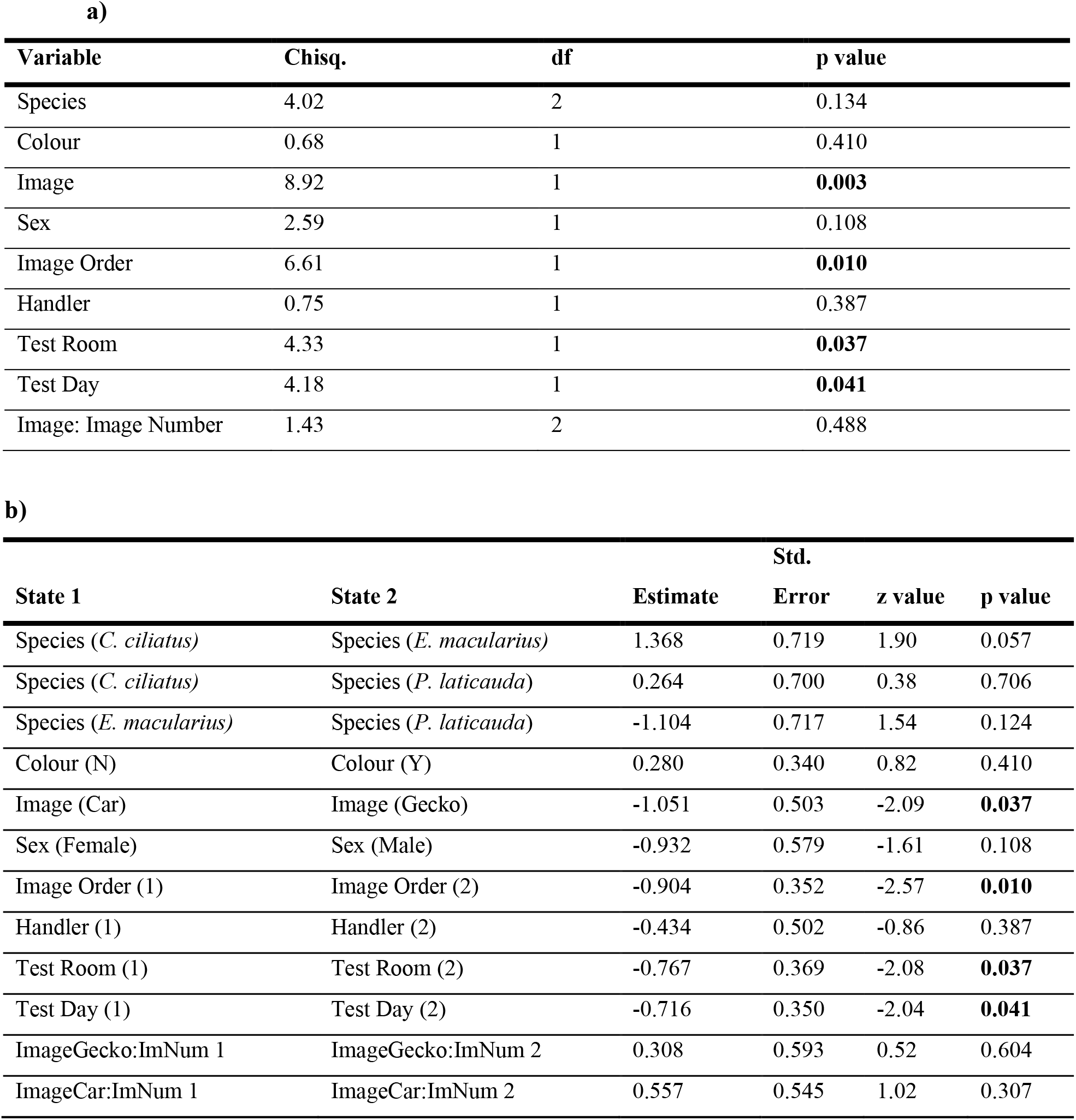
Summary output from a mixed effects logistic regression model on the reaction of geckos to images. **a)** Chi-square tests on effects **b)** Wald tests on parameters. The results show state 2 in comparison to state 1. A negative estimate signifies that the state 2 is less likely to respond than state 1 (e.g., a gecko is less likely to respond to Image (Gecko) (state 2) when compared against Image (Car) (state 1). Significant p-values (<0.05) are highlighted in bold.

We found that geckos responded significantly less to images of geckos than cars (p=0.037) – independently of colouration –, to the second image than to the first (p=0.010), to the second test room than to the first (p=0.037), and during the second day of the experiment than to the first (p=0.041) (Table 2b). We found no differences in responses to the two different images with the same content (p*=*0.307 for the car images and p*=*0.604 for the gecko images; Table 2b). When looking at the interaction among variables, we observed that, depending on the image type (gecko or car), animals responded differently to the image order (p=0.007); post-hoc pairwise comparisons indicate that animals responded significantly less to the second image compared to the first image in a single test day when the image is a gecko (p=0.002), but not when the image is a car (p=0.85). We found no significant interaction among all the other tested variables (p>0.05). Although a different response was observed between the two experimental rooms, there were no significant interactions with other variables, and thus the pattern of responses did not differ between rooms. When we analysed data for *touching*>0 only, we found that only the image order had a marginally significant effect (p=0.04).

### 3.3 Influence of factors on duration of the response (PLT>0)

In addition to analysing the variables affecting the presence or absence of a response, we also tested which variable(s) influenced the proportion of time geckos spent reacting to the images (*PLT*>0; Table 3). The generalized linear mixed effects model using a gamma distribution indicates that species responded differently (p<0.001). None of the other variables – including colour information – influenced how long a gecko responded to images (Table 3a). When looking at the influence of multiple levels within each variable, we found that the diurnal species, *P. laticauda*, spent more time interacting with the images than the two nocturnal species (p<0.001 in both cases; Table 3b). No significant interaction effects were observed (p>0.05). When we analysed data for *touching* only, *P. laticauda* also spent significantly more time *touching* the images than the other two species (p=0.002).

**Table 3.** Summary output of the generalized linear mixed effects model based only on the animals that responded to the image (PLT>0). **a)** Chi-square tests on effects. **b)** Wald tests on parameters. The results show state 2 in comparison to state 1. A negative estimate signifies that the state 2 is less likely to respond than state 1 (e.g., a gecko is more likely to respond if they are sex (Male) (state 2) when compared to Sex (Female) (state 1)). Significant p-values (<0.05) are highlighted in bold.

**a)**
| Variable | Chisq. | df | p value |
| --- | --- | --- | --- |
| Species | 25.44 | 2 | **<0.001** |
| Colour | 0.02 | 1 | 0.90 |
| Image | 0.03 | 1 | 0.87 |
| Sex | 3.84 | 1 | 0.05 |
| Image Order | 2.32 | 1 | 0.13 |
| Handler | 0.91 | 1 | 0.34 |
| Test Room | 0.16 | 1 | 0.69 |
| Test Day | 0.03 | 1 | 0.85 |
| Image: Image Number | 3.75 | 2 | 0.15 |

**b)**
| State 1 | State 2 | Estimate | Std. Error | t value | p value |
| --- | --- | --- | --- | --- | --- |
| Species (*C. ciliatus*) | Species (*E. macularius*) | 0.999 | 2.220 | 0.45 | 0.652 |
| Species (*C. ciliatus*) | Species (*P. laticauda*) | 15.371 | 3.199 | 4.81 | **<<0.001** |
| Species (*E. macularius*) | Species (*P. laticauda*) | 14.372 | 3.105 | 4.63 | **<<0.001** |
| Colour (N) | Colour (Y) | -1.65 | 1.270 | -0.13 | 0.897 |
| Image (Car) | Image (Gecko) | 1.381 | 2.239 | 0.62 | 0.5375 |
| Sex (Female) | Sex (Male) | 4.151 | 2.119 | 1.96 | 0.050 |
| Image Order (1) | Image Order (2) | 2.004 | 1.317 | 1.52 | 0.128 |
| Handler (1) | Handler (2) | 1.737 | 1.820 | 0.95 | 0.340 |
| Test Room (1) | Test Room (2) | 0.545 | 1.350 | 0.40 | 0.686 |
| Test Day (1) | Test Day (2) | -0.242 | 1.305 | -0.19 | 0.853 |
| ImageCar:ImNum 1 | ImageCar:ImNum 2 | -1.415 | 1.942 | -0.73 | 0.466 |
| ImageGecko:ImNum 1 | ImageGecko:ImNum 2 | -3.935 | 2.337 | -1.68 | 0.092 |

## Discussion

Our study focused on understanding whether colour information impacts the perception of a biologically relevant and familiar object vs. a biologically irrelevant and novel object using 2D images. Previous research has demonstrated that some lizard species perceive videos of conspecifics as conspecifics (e.g., Macedonia, Evans & Losos, 1994; Ord *et al*., 2002, Frohnwieser *et al*., 2017), suggesting that the ability to recognize two-dimensional images as real objects might be widespread across lizards. We therefore expected geckos to perceive and interact with the 2D images shown to them. We also expected that geckos would interact more often and longer with the coloured image than with the one shown in grayscale for the familiar object (an image of the gecko), because of its biological significance, but we did not expect to find a difference for the novel unfamiliar object (an image of the car).

Our results revealed that the geckos showed different responses depending on the image content – car or gecko (Table 2). Geckos and responded more frequently to the novel images (car) than to an image of a conspecific gecko, independently of colouration (Table 2). To our knowledge, this is the first study that has examined the effect of novel images on the behaviour of geckos, but studies in mammals, fish, and other reptiles have found similar results, where more time was spent interacting with novel objects (e.g., Antunes & Biala, 2012; Lucon-Xiccato & Dadda, 2016; Nagabaskaran *et al*., submitted). The lower response observed for the gecko image in our study may be due to the image lacking other sensory stimuli associated with a “real” object (e.g., movement, pheromones, visual displays, three-dimensional orientation, UV reflectance) and commonly used in individual or species recognition (d’Eath, 1998; Bovet & Vauclair, 2000; Ord *et al*., 2002; Mason & Parker, 2010).

In our study, geckos did not rely on chromatic information to perceive and discriminate between images, even though these animals are known to be tetrachromatic (Röll, 2000; Roth & Kelber, 2004). In fact, we observed that while geckos discriminate the type of image, they did not react differently if this was in colour or grayscale, independently of the biological relevance of the image (Table 2). The observed lack of support for a role of colour information as a visual stimulus was surprising, since the importance of colour vision and coloured traits in lizards are well established (e.g., LeBas & Marshall, 2000; Klomp *et al*., 2017; Ossip-Drahos *et al*., 2018; Kabir *et al*., 2019; Allen *et al*., 2020; Dollion *et al*., 2020). It is to note that in our work we tested for chromatic differences in colouration (hue and saturation) and not achromatic ones (brightness), which may be an important component of colouration. However, the role of colouration in improving object perception is not clear (e.g., Delorme *et al*., 2000, Tanaka *et al*., 2001), and our findings support the only other paper that has investigated this in reptiles (Frohnwieser *et al*., 2017). Colour may act in combination with shape information, or shape may be the main driver of discrimination, suggesting a more limited role of colouration in object perception (Tanaka *et al*., 2001; though see Frohnwieser *et al*., 2017). Our results seem to suggest that geckos discriminate between the two 2D stimuli independently of colour information, therefore shape differences may be more important for the perception of the object than colouration. Shape is considered to be important in conspecific recognition and there is evidence to suggest that other species can discriminate between conspecifics and objects that are shaped similarly to them (Johnson & Horn, 1988; Palmer, Calvé &Adamo, 2006). Although in our work shape seems to be more important than colour for object perception, the use of two-dimensional printed images might not adequately reflect some properties related to colour vision, such as pigmentation or structure of the colour or reflectance in the UV spectrum of the real object that is being tested (e.g., D’Eath, 1998). In our work, the lack of increased response to coloured images, especially for the gecko images, may also be due to the fact that colour may be used in combination with other sensory stimuli such as movement, UV reflectance, or scent, which are known to play a big role in response to conspecifics in lizards (e.g., LaDage *et al*., 2006, Kabir *et al*., 2019; though see Frohnwieser *et al*., 2017).

We found that within the same testing day, geckos reacted less to the second image of a gecko than the first, regardless of whether the colour/grayscale images were shown first. This suggests that geckos might become habituated to an image if the image type has social significance (e.g., another gecko). The same habituation was not observed when looking at the car image during the same day. However, when looking at response to the same image type over two consecutive days, we observed a lower response during the second day of testing independent of the image content (Table 2). Nevertheless, a lower response was not observed when another image of the same image type was shown after six weeks (Table 2). This suggests that the lizards habituated to the stimulus types after short-term exposure and did so more rapidly for the gecko than the car. However, this habituation effect was not observed after a more substantial gap. This fits with what we would expect to see from work with mammals and birds (Shettleworth, 2009) and suggests that this sort of paradigm could be a useful approach to testing learning and retention in this group.

The visual components that are important to an animal after noticing an object might be different from the factors that initially drew their attention to the object, and positive responses were therefore analysed separately. We found that when analysing only the positive responses (*PLT*>0), there was an overall lack of distinction in time spent interacting with distinct image types, which suggests that image content may be more important for distinguishing between image types, but other aspects of a stimulus may be important for prolonged interaction. When analysing only *PLT>0*, we found that *P. laticauda* responded significantly more than the other two species (Table 3). This distinction could be due to different responses to visual stimuli between nocturnal and diurnal species. Although nocturnal species, such as *C. ciliatus* and *E. macularius* are able to see colour in dim light (Kelber & Roth, 2006), it is currently not known what they primarily use vision for. Previous studies on diurnal geckos have found that they can perceive and discriminate between different colours and coloured features of conspecifics (Ellingson, Fleishman & Loew, 1995; Hansen *et al*., 2006; Minnaar *et al*., 2013), suggesting that these species may rely on vision more than nocturnal ones. Future studies including other diurnal and nocturnal species could further test this hypothesis.

Our work supports the capacity of geckos to perceive and discriminate among images with different content, and that colour information may be less important in improving this discrimination in the absence of other sensory stimuli. Our study also suggests that while all the three tested species respond to visual stimuli, diurnal geckos such a *P. laticauda* may rely more on vision for object processing than nocturnal species. Future studies should aim to further investigate the role of colouration for object perception and discrimination in combination with other sensory information and the influence of acuity and sensitivity to different colours and colour components in geckos, as this will have implications on how colour is used in these animals.

## Data Availability

Data will be provided as supplementary material after manuscript acceptance

## Acknowledgments

We would like to thank Nickolas Moreno, Spencer Potter, Hannah Altonji, Blake Meador, Onree Wilson, and Trevor Rayl for help with gecko husbandry and creating the enclosures for the experiments. We are thankful to Yulia Bereshpolova for discussion about vertebrate vision in the early stages of this work. We are grateful to Aamod Zambre for providing critical comments on this manuscript. This work was funded by a Gulf Coast Advance Fellowship and by the University of South Alabama Research Development Funds to YC.

## Author Contributions

YC, NK, AW, MK, SG, conceived the project and designed the experiments. NK and MR performed the experiments. BW, JC, and YC performed the statistical analyses. YC and NK wrote the manuscript, SG and AW provided critical revision, and JC, MK, BW provided comments. YC obtained funding for this project. All authors have read and approved the final manuscript.

## Supplementary Material Figure Legend

**Figure S1**. Experimental set up showing the testing arena, the placement of the image within the arena, and the location of the video-cameras outside of the experimental arena (*a* and *b* images). Testing arena shown in the images did not have the three sides covered with white paper (see Methods section).

**Figure S2**. Images of the geckos and cars used for each species and for each dataset. For each image type, the two image numbers are shown.

**Figure S3**. Protocol used for habituation and experimental set up.

**Figure S4**. Example of behaviours analysed from the experiments. A) *Touching*: When a gecko’s tongue or the snout was touching the image. B) *Looking*: When a gecko’s head and eyes were directed directly towards the image. C) *Not Looking*: When a gecko’s head is not directed at the image (see Supplementary Information for additional details).

**Figure S5:** Frequency (top) and distribution (bottom) of PLT on log transformed (top) and raw (bottom) data for each species.

