## Supplementary Material for "Does colour impact attention towards 2D images in geckos?"

***Detailed Materials and Methods***

***Study Subjects and Captivity Conditions***

All geckos were housed in the same room at the University of South Alabama and exposed to 12-12 hour light-dark cycles, with the individuals of *P. laticauda* having access to additional UV lamps (ReptiSun 10 UVB bulbs), as required by the biology of this species. The containers for *E. macularius* and *C. ciliatus* were plastic boxes (*E. macularius*: 58 cm L x 41 cm W x 15 cm H; *C. ciliatus*: 46 cm x 31 cm x 17 cm for males and 43 cm x 28 cm x 32 cm for females) with newspaper bedding. The *P. laticauda* were in glass cages (Exoterra, 30 cm x 30 cm x 45 cm) with cocoa mulch bedding. Housing conditions were different among species according to different species requirements. The room temperature was kept stable between 22-25 °C and each enclosure contained a moist hiding place or fake plant and a heat pad for thermoregulation. Humidity level and temperature in the room were checked daily. The health condition of each individual was also checked daily. No geckos were tested during shed. Geckos were fed three times a week with a combination of crickets dusted with calcium and vitamins, mealworms, and a fruit supplement. Feeding schedules were not changed for the geckos involved in the experiment.

***Experimental Set-Up***

Each gecko was given a unique individual code for the experiments and could be identified by physical characteristics, such as colour pattern, size, and cage number. Experiments were carried out concurrently in two separate test rooms with identical setups. Although the experimental setup in the two rooms was the same and the rooms were similar in size with no external sources of noise or light, each room contained different furniture, floor (e.g., one room has carpet and the other has bare floor) and items that could not be removed. However, since the experimental arena had its sides covered by white sheets, geckos could not see any differences in the surrounding during the experiments. Experimental rooms were accessed only by the testers (NK and MR) when needed and were not used before and during the entire length of the experiments. Since we could not account for smell or other features of each room undetectable to humans, test room was used as a fixed effect in our analyses and all the individuals and all the images were tested in both rooms. We observed a different interaction rate (called also response here) of the geckos between the two rooms (see Results). The lower response rate obtained in one of the room was however across experiments and not associated with any of the other tested variables (no interaction effects were observed among variables). Difference in response between the two rooms may be due to smell or other features undetectable to any of us.

For the diurnal species, the overhead lights of the room (GE T8 Starcoat ECO bulbs, 32W) were kept on throughout all portions of the experiment. For the nocturnal animals, the overhead lights of the room were turned off and one lantern (Kino Flo CFL 26W KF55 lightbulb) was hung about 1.5 m off the ground and 3 m away from the testing arena to provide ~4-7 lux depending on where the light was measured in the rooms. Illuminance was measure in different points of the rooms using the phone app *Light Meter* on a Samsung Galaxy Note 8. The different light set-up between nocturnal and diurnal species was used to better reflect the distinct time of activity – and captivity conditions – between nocturnal and diurnal animals, as diurnal and nocturnal animals are known to perform best when light conditions reflect those of their main activity time (Higham and Schmitz 2019).

The testing arena (30.5 cm x 61 cm x 20.3 cm) for the experiments was constructed out of clear plastic and silicone. The lid of the arena was just large enough to wrap around each side and was secured by clear tape in order to prevent geckos from escaping. The size of the arena ensured that the images shown for the experiment fitted within the narrow side of the testing arena and enough room was provided for each tested gecko to move and react freely. The other three sides and the bottom of the arena were covered in white tissue paper placed on the outside of the arena and never in contact with the animals. One narrow side of the arena and lid were not covered by paper to allow images to be shown and to capture behaviours using the video cameras placed outside and above the arena. The uncovered side where the image was shown was always facing the same direction in each room and the only objects visible beyond the transparent images (see below) were a white wall and the tripod stand. In order to record the experiments, two video cameras (Panasonic HC-V770 and Canon Vixia HFR700) were set on tripods and situated at each end of the arena. These video cameras were chosen because of their characteristic of recording good quality videos in dim lighting.

***Test Images***

The images of the geckos used for experiments were obtained by taking pictures of two randomly selected individuals among the ones we have for each species. In order to ensure that all geckos had the same images to respond to, some individuals were shown the photographs of themselves during the experiment. The geckos for which their own image was shown did not respond differently from the other geckos of the same species (data not shown). Pictures were taken using a DSLR camera (Canon rebel t6i) on automatic settings in the experimental room with only the overhead lights of the room being used and the gecko in the testing arena. All of the geckos were photographed from the same side of each respective animal (Supplementary Material Fig. S2). All pictures were taken showing the lateral view of the animals or the cars and were oriented similarly. The lateral view was used to simulate what an animal would see if it were on the same plane of the stimulus. Although two of the species used could climb and therefore look at the images from different angles, we decided to keep the plane of view similar among the tested species in order to show the same stimulus to different receivers. For the car, images were obtained from the web using a Google image search and selected based on the quality of the image and clear colour and colour pattern similar to the one of the species tested. Before being printed, all gecko and car images were scaled to actual animal size, so that the size of the object shown was not discriminative between car and gecko images.

Pictures were printed on transparent sheets (8.5" x 11") in order to show the image without confounding effects of the background as geckos could respond to a background effect more than to the image itself. To test if the geckos responded differently if the image is shown in colour or not, for each image type, we used both true natural colour of the object (gecko or car) and grayscale. Although geckos are known to be able to see in the UV spectrum, in our experiments we wanted to focus only on the comparison between colour vs. non colour, without the additional information provided by UV colouration or different components of the colour itself (hue, brightness, etc. were not manipulated). Grayscale was used as an alternative to natural colour to eliminate the hue and saturation and leave just the brightness and intensity of each pixel in a given image. Grayscale images were converted from the colour image using GIMP's image filtering tools by using the pre-programmed grayscale command in the "image" menu bar. The grayscale conversion function uses the weighted sum of the red, green, and blue components of the coloured image and transforms them into a single-channel grayscale image. Each image type for each individual and in each colour was printed for each tested individual in order to avoid that scent from a previously tested animal could be left on the image and influence the response of the animal tested after. A total of 368 images were printed in total.

***Testing Procedure***

Experiments were run between October 2017 and May 2018. The length of the habituation period at the beginning of each experimental day was chosen based on preliminary trials carried out during summer 2017 in which geckos felt comfortable in the testing environment (i.e., eating, drinking, exploring) by three days. All geckos were habituated to the arena for three consecutive days before beginning the experiments (Supplementary Material Fig. S3). After the three-day habituation period, the geckos began the testing phase of the experiment. Geckos were tested individually and were placed in their own testing arena for three consecutive hours for each day of habituation (Days 1-3). After each daily habituation session, the gecko was moved back to its normal housing terrarium and was not used again until the following day. Diurnal geckos were habituated between 8am-5pm and nocturnal geckos were habituated between 5pm-11pm each day. These times were chosen to more accurately reflect the period that each species is most active in their natural habitat (i.e., diurnal vs nocturnal activity). Once in the testing arena, the geckos were supplied with a plastic dish filled with water and a single cricket during each habituation day. Geckos were considered to have habituated and be comfortable with the testing arena when they show a normal behaviour (eat or drink) at any time during habituation. Normal feeding schedule was not changed for the tested individuals during the experiments, so the cricket provided during habituation was additional to the food normally provided to the animals. During the habituation time, three sides of the testing areas were covered with a white tissue paper and the remaining side was covered with a removable sheet of white printer paper that was used to cover the image later in the experiment (see below). The test images were not included in the habituation setup to ensure that the animal was not able to see the image before the test.

Each experiment started with an acclimation period. From preliminary test trials, ten minutes was enough time to gauge an animal's behaviour before it becomes disinterested and no longer interacts with the image. After the two pictures (Supplementary Material Fig. S3) were shown on each experimental day, the animal was removed from the testing arena and the arena was prepped for the next tested animal by washing it with soap and water to remove scents or debris. Throughout the experiment, only two researchers, NK and MR, handled the geckos during the trials. Ultimately, the animal was left alone in the room during all phases of the experiment besides the initial handling, the removal of the cover sheet, the changing of the image, and the conclusion of the experiment. On the following day, the order of the images shown (called "image order") was reversed (i.e., from colour/grayscale to grayscale/colour) in order to avoid the geckos responding to the order of the images rather than the colours (Supplementary Material Fig. S3). Only a single image type (e.g., car or gecko) was tested during each 5 days trial in order to minimize the effects of image habituation or memory on responses, and the same gecko was not presented with any other image for about three weeks (Supplementary Material Fig. S3).

Image order for each gecko tested was as follows: (1) a single gecko image was shown in colour and grayscale during the first week; (2) three weeks later, a single large car image was shown in colour and grayscale; (3) six weeks after seeing the first gecko image, the second gecko image was shown in colour and grayscale; (4) finally, six weeks after seeing first large car image, the second large car image was shown in colour and grayscale (Supplementary Material Fig. S3). Thus, for each gecko tested, each dataset required approximately nine weeks of testing. The break of about six weeks between showing different images within the same image type (e.g., two gecko individuals) was chosen to decrease the chance that animals would anticipate the image type to be observed based on familiarity of image order. In our experimental set up, we therefore tested for differences in responses based on species, sex, image colour, image type, different images of the same type, image order based on colour, as well as habituation or memory to the experimental setup.

### *Data Collection*

In order to collect data, we used video footage from both cameras to measure the amount of time each gecko was interacting and looking at the images for each experiment. The categories that were used for the analyses were *touching*, *actively looking*, *looking-inactive*, and *not looking*. The behaviours observed in this study were easily measurable reactions that were observed in our test trials (Summer 2017), and were both easily quantifiable and consistently seen between individuals and among species. The geckos were considered to be interacting with the image if after being able to see it, they actively looked at the image or touched the image directly. To assess if the gecko was *actively* versus *inactively* looking, we used the amount of time since the animal last moved to make the distinction. Based on pilot experiments, if the gecko did not move for at least ten seconds then they were considered as *inactive* for that time. Since geckos can have periods of inactivity followed by quick movements, ten seconds was selected as the cut-off in order to ensure that the lack of movement was not just a momentary stop. Although during the period considered *inactive* a gecko could potentially still process visual images, since there is no way to assess this without using invasive methods our approach is more conservative in not over-estimating the time of interaction with an image. A gecko was considered to be *touching* when its nose, tongue (licking) or snout were in contact with the image. The difference between *looking* and *not looking* depended on if the gecko's head was directed towards any part of the image or if it was turned elsewhere. Geckos exhibit binocular vision, although it is not known if it is stereoscopic (Röll 2001); using only the time where the gecko appears to focus both eye on the image, rather than including time where the image might be seen in their peripheral, makes for a more conservative estimation of responsiveness to the image. Instances in which geckos never turned towards the image or had the ability to see it were removed from the analysis.

***Supplementary Material Table S1: Description of the variables used in this study.***

| **Variable** | **Explanation** |
|---|---|
| Species | What species was being tested (*E. macularius*, *C. ciliatus*, and *P. laticauda*) |
| Sex | The sex of the gecko being tested. |
| Image Type | Differentiates between gecko and car images. |
| Colour | Whether the image was in colour or grayscale |
| Image Number | It refers to which of the two images of the same type (e.g., gecko or car) was shown for each image type. Image number was always analysed depending on Image Type. |
| Image Order | Denotes which image (colour or grayscale) of the same image type was shown first and second during a single testing day. All the images were shown in both colour and grayscale as first or second across all the experiments. |
| Room | Designates which room a gecko was tested in on a given test day. |
| Handler | Which researchers handled the geckos and conducted an experiment in a testing day. |

| | |
|---|---|
| Day | The day in which we ran the experiment within the same week |

***Supplementary Material Table S2: Summary statistics of PLT measure by species.*** "Species" refers to the species of gecko being tested. "N" indicates the total number of PLT measures, while "NA" refers to the number of missing values (gecko never saw the images), "$N_0$" refers to the number of measures with PLT=0, and "$N_1$" refers to number of measures with PLT>0. "Max" shows the maximum PLT measures for each of the three species. "Median" and "SD" shows the median and standard deviation of PLT, respectively. "Shape" and "Ratio" are two estimated parameters of the Gamma distribution fitted to PLT>0.

| **Species** | **N** | **NA** | **$N_0$** | **$N_1$** | **Max** | **Median** | **SD** | **Shape** | **Ratio** |
|---|---|---|---|---|---|---|---|---|---|
| *C. ciliatus* | 124 | 35 | 22 | 67 | 0.304 | 0.053 | 0.069 | 1.562 | 18.866 |
| *E. macularius* | 128 | 10 | 14 | 104 | 0.311 | 0.066 | 0.056 | 1.928 | 25.176 |
| *P. laticauda* | 97 | 10 | 21 | 66 | 0.133 | 0.033 | 0.029 | 1.422 | 38.543 |
