## Supplementary figures and images for "Does colour impact attention towards 2D images in geckos?"

### Supplementary Material Fig. 1

(a)

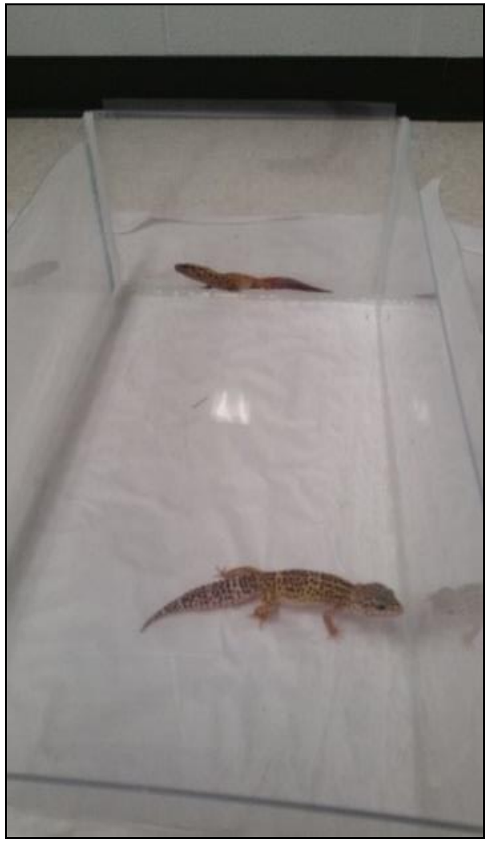

(b)

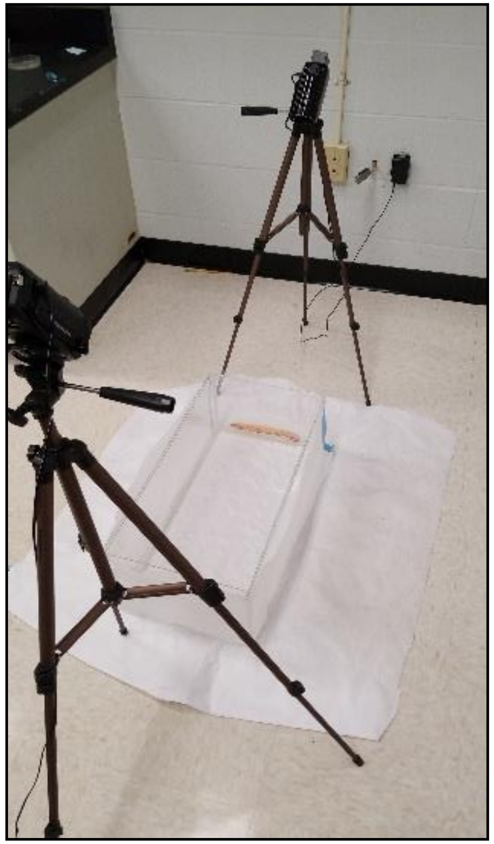

### Supplementary Material Fig. 3

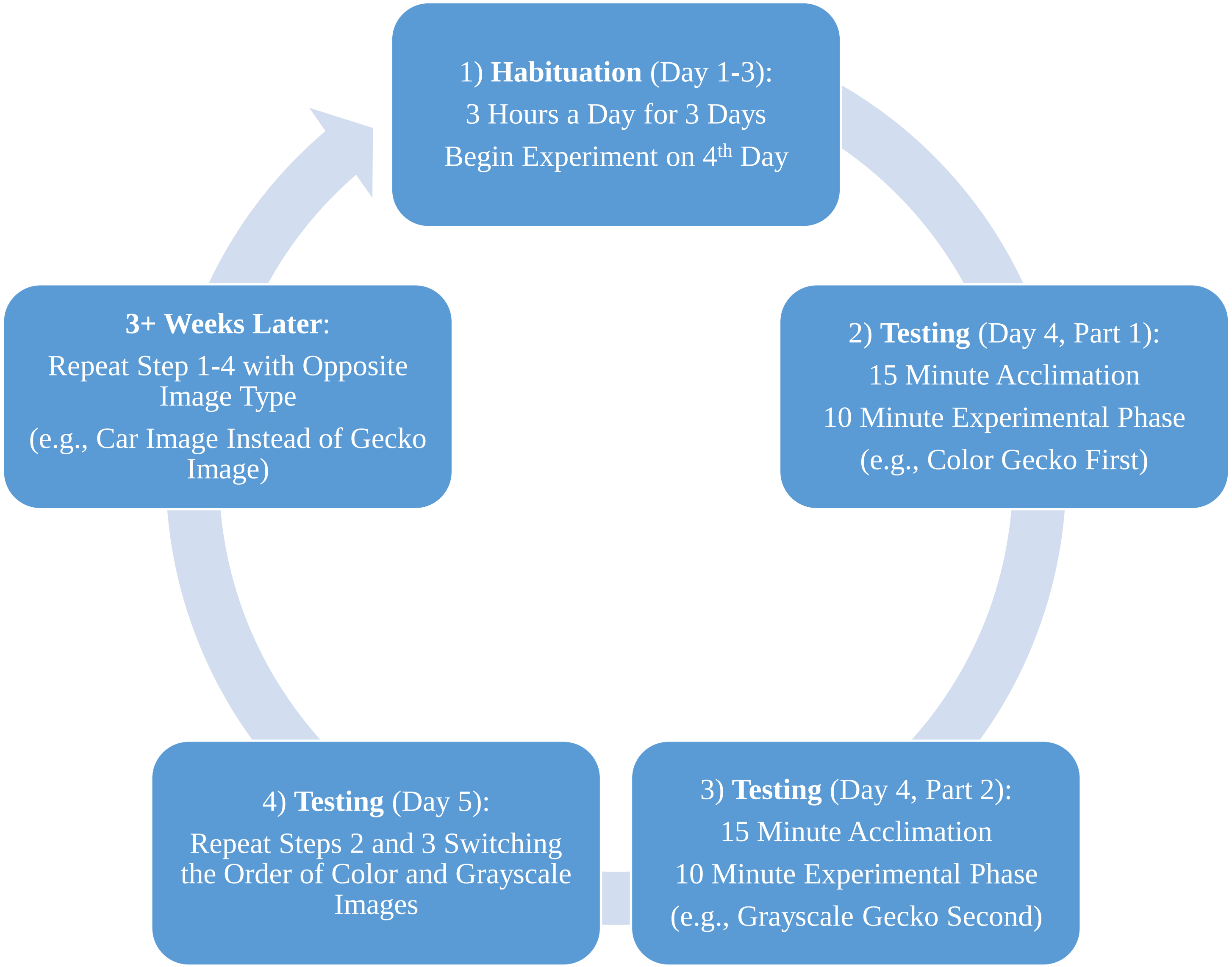

### Supplementary Material Fig. 4

(a)

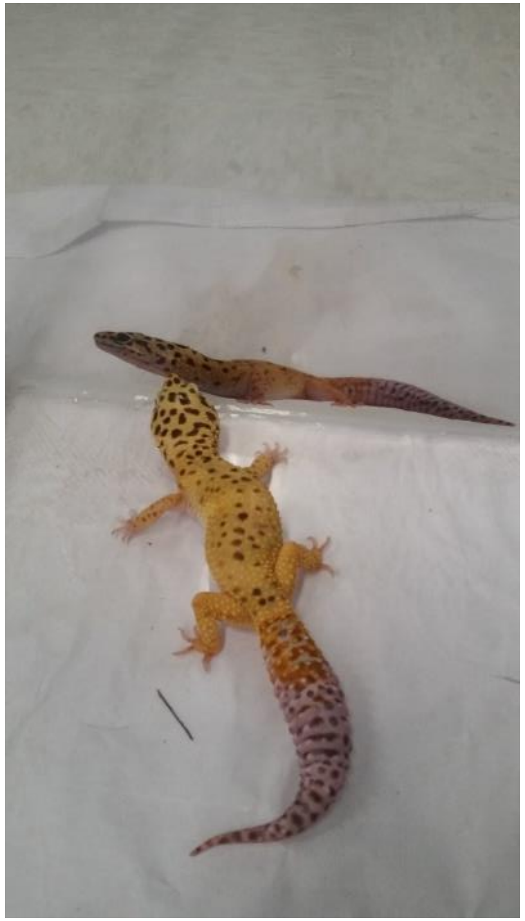

(b)

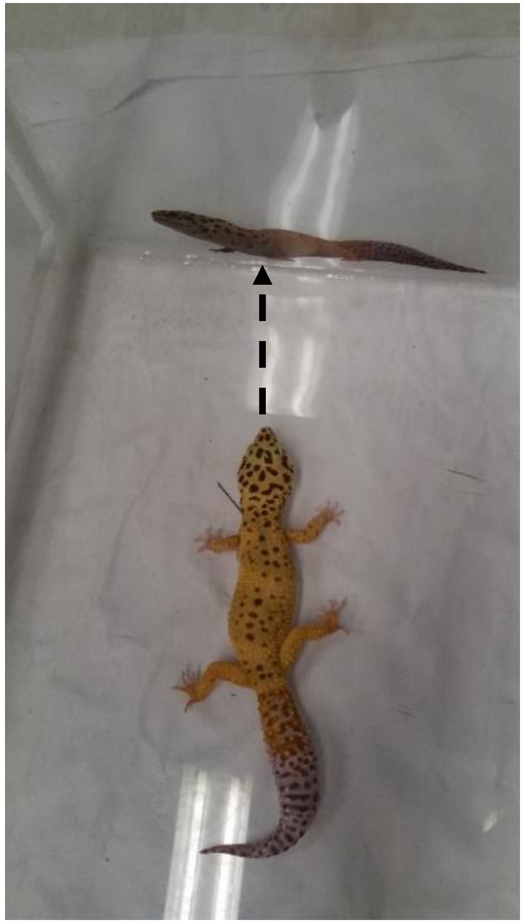

(c)

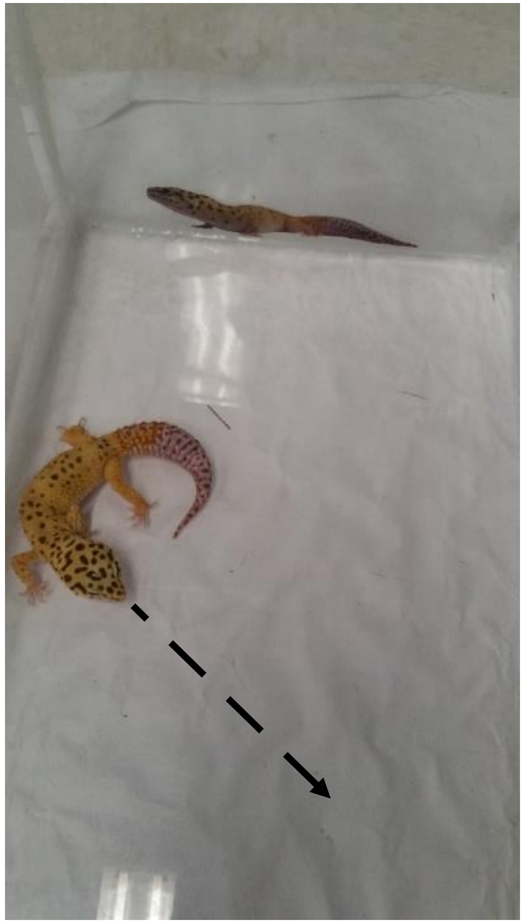

### Supplementary Material Fig. 5

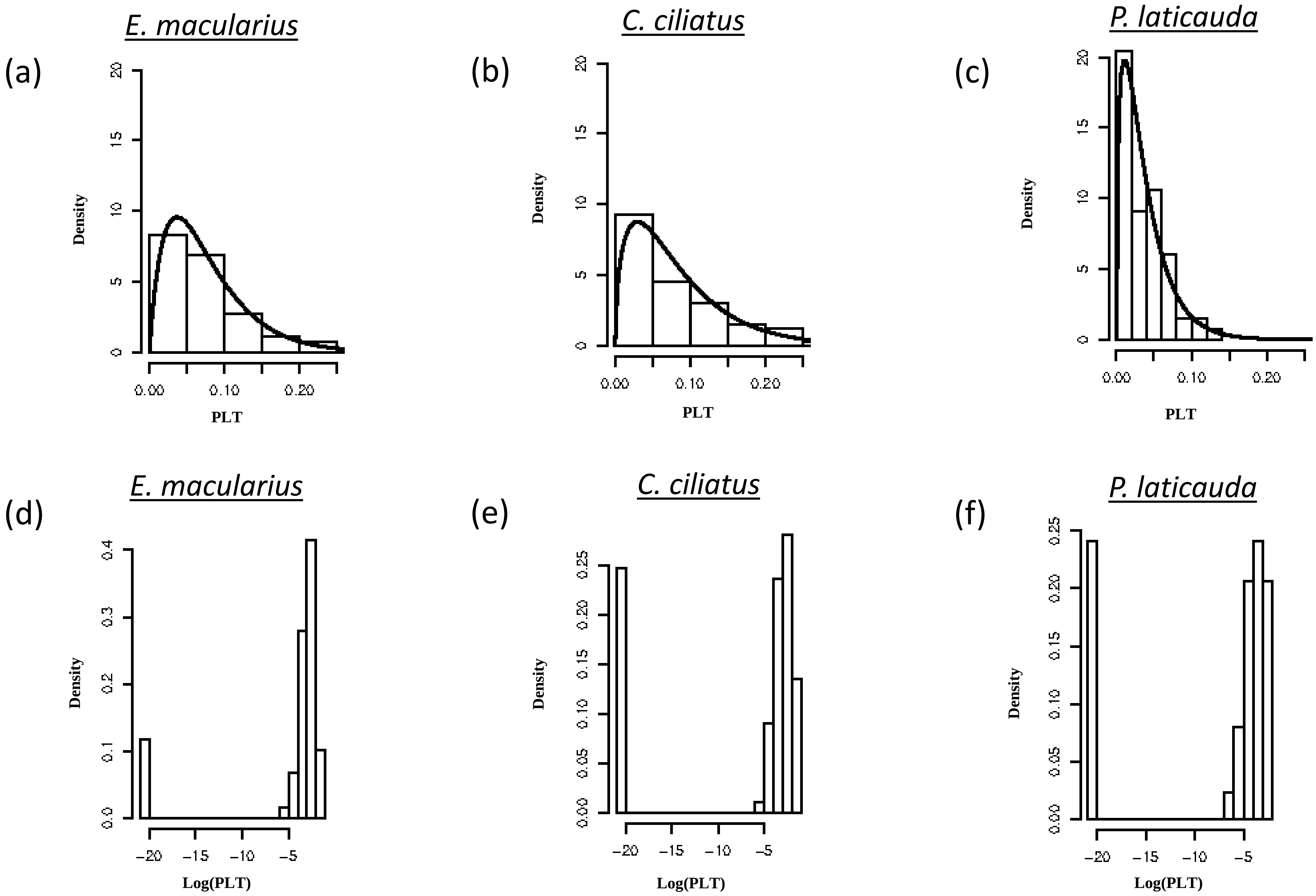
