## Supplementary Material Fig. 2 for "Does colour impact attention towards 2D images in geckos?"

| | *E. macularius* Image 1 | *E. macularius* Image 2 | *C. ciliatus* Image 1 | *C. ciliatus* Image 2 | *P. laticauda* Image 1 | *P. laticauda* Image 2 |
|---|---|---|---|---|---|---|
| **Gecko Color** | 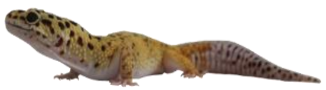 | 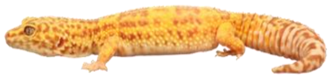 | 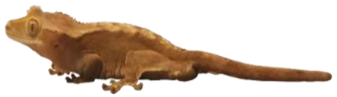 | 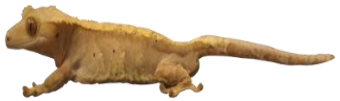 | 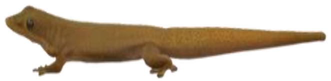 | 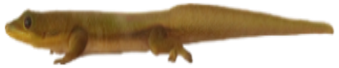 |
| **Gecko Gray** | 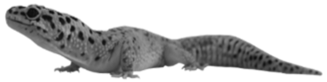 | 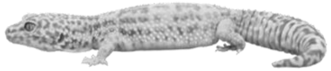 | 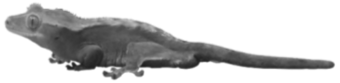 | 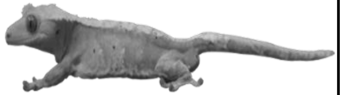 | 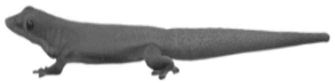 | 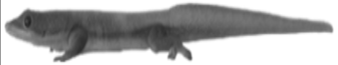 |
| **Car Color** | 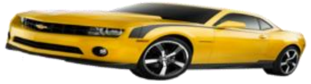 | 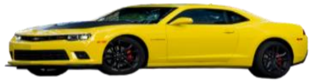 | 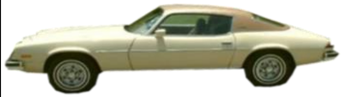 | 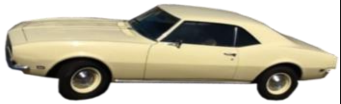 | 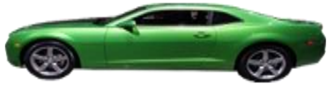 | 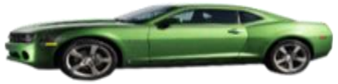 |
| **Car Gray** | 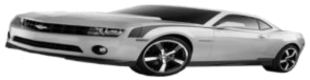 | 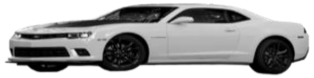 | 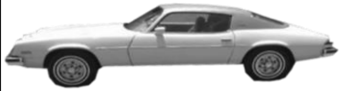 | 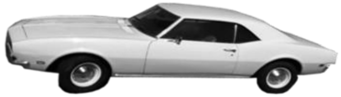 | 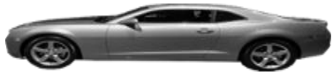 | 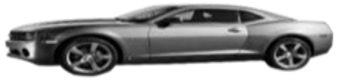 |
